# The tuberculosis-associated microenvironment promotes HIV-1 persistence by impairing CD8+ T cell-mediated viral control

**DOI:** 10.1101/2025.10.14.682453

**Authors:** Samantha Cronin, Jennifer Simpson, Andrea Pereyra Casanova, Yuchen Li, Josefina Marín-Rojas, Freja Warner van Dijk, Katie Fisher, Zoï Vahlas, Thomas O’Neil, Kirstie Bertram, Eunok Lee, Gabriela Turk, Florencia Quiroga, Anthony Kelleher, Christel Vèrollet, Luciana Balboa, Sarah Palmer, Gabriel Duette

## Abstract

*Mycobacterium tuberculosis* (*Mtb*), the causative agent of tuberculosis (TB), is the most common coinfection in people living with HIV-1 (PLWH). This coinfection is associated with accelerated HIV-1 disease progression and reduced survival. However, the immunological and virological mechanisms driving this progression are incompletely understood. To address this knowledge gap, using pleural effusion samples from PLWH and TB, we investigated the HIV-1 genetic landscape and the anti-HIV-1 immune response impacted by a TB-associated microenvironment. Our results revealed an enrichment of genetically intact HIV-1 and impaired CD8+ T cell-mediated antiviral response at the site of HIV-1/*Mtb* coinfection. These findings indicate that the TB-associated microenvironment promotes the persistence of cells infected with replication-competent HIV-1 by creating a niche of reduced antiviral immune pressure, potentially contributing to the worsened clinical outcomes observed in PLWH and TB.

## Introduction

Despite considerable progress with antiretroviral therapy (ART) improving the health and life expectancy of people living with HIV-1 (PLWH), an effective cure remains elusive. The persistence of a reservoir of memory CD4+ T cells infected with genetically intact and replication-competent HIV-1 is a major barrier to eradication. Although the majority of proviruses in PLWH are genetically defective (1), even a small pool of replication-competent viruses is sufficient to drive viral rebound upon treatment interruption or failure (2-4). A deeper understanding of the factors shaping the size and genetic features of the persistent HIV-1 reservoir is essential for the development of new, effective curative strategies.

Several immune mechanisms contribute to HIV-1 persistence. Latency and proliferation of HIV-1-infected cells are central drivers of reservoir maintenance despite effective ART (5, 6). Additionally, although CD8+ T cells are critical for controlling HIV-1 during acute infection, they eventually acquire an exhausted phenotype that undermines viral control (7, 8). In addition to chronic activation and exhaustion, tissue-resident CD8+ T cells, exhibit reduced cytotoxic capacity, allowing the virus to persist within anatomical niches such as lymph nodes (9, 10). Thus, inefficient CD8+ T cell-mediated antiviral activity allows for the persistence of genetically intact HIV-1 which contributes to viral rebound when ART is interrupted.

*Mycobacterium tuberculosis (Mtb),* the causative agent of tuberculosis (TB), is the most common coinfecting pathogen among PLWH. TB remains a major global health threat, responsible for 10.8 million new infections and 1.25 million deaths annually (11). HIV-1/*Mtb* coinfection has been described as a *syndemic*, with the concurrence of both pathogens exacerbating the clinical severity of each other (12, 13). PLWH with concurrent TB are 18 times more likely to develop active TB and face reduced survival compared to individuals with one of these infections (14, 15). Consequently, TB is the leading cause of death among PLWH, accounting for 33% of AIDS-related deaths in 2023 (11, 16).

Despite its frequency and clinical severity, little is known about how TB influences the size and genetic composition of the replication-competent HIV-1 reservoir. Studies of blood-derived cells have yielded conflicting results, with some reporting increased HIV-1 DNA during TB coinfection and others reporting decreases (17-19). This inconsistency highlights the need to investigate the systemic and compartment-specific effects of HIV-1/TB concurrence. While *Mtb*-specific CD4+ and CD8+ T cell responses are known to be impaired during TB disease, whether TB also compromises HIV-1-specific CD8+ T cell responses, potentially allowing for the pool of HIV-1-infected CD4+ T cells to expand, remains unexplored (20).

Pleural effusion (PE), an accumulation of fluid in the pleural space, occurs in approximately 30% of TB patients (21). It results in response to the presence of mycobacteria or mycobacterial antigen in the pleura, triggering local inflammation and leukocyte infiltration (22, 23). In PLWH/TB, PE is more common and contains higher HIV-1 titres than paired plasma (24-26). Thus, TB-PE represents a physiologically relevant *ex vivo* model to study the microenvironment at the site of HIV-1/*Mtb* coinfection and its impact on HIV-1 persistence (27-30).

In this study, we genetically characterized HIV-1 from TB-PE and examined the impact of TB-PE on antiviral CD8+ T cell responses. Our findings suggest that the TB-associated microenvironment creates a niche of reduced anti-HIV-1 immune pressure, permissive to the persistence of replication-competent HIV-1, which may contribute to the worsened clinical outcomes observed in PLWH/TB.

## Results

### HIV-1 proviral genetic landscape in people living with HIV-1 and TB

Despite the clinical consequences of HIV-1/*Mtb* coinfection, the impact of active TB on the HIV-1 proviral genetic landscape within circulating HIV-1-infected cells remains unclear. To address this, we analyzed HIV-1 proviral sequences from peripheral blood mononuclear cell (PBMC) samples obtained from five people living with HIV-1 (PLWH) and five individuals with HIV-1 and active TB (PLWH/TB) (Table 1). The duration of ART ranged from 1 to 4 months for most participants, except for one individual who had been on ART for eight years and experienced treatment failure (Table 1). Viral loads ranged from <50 to 1,180 copies/ml, with no significant differences observed between the two groups (Figure 1A). Consistent with previous studies, CD4+ T cell counts were significantly lower in PLWH/TB (Figure 1A)(31, 32).

**Figure 1.**
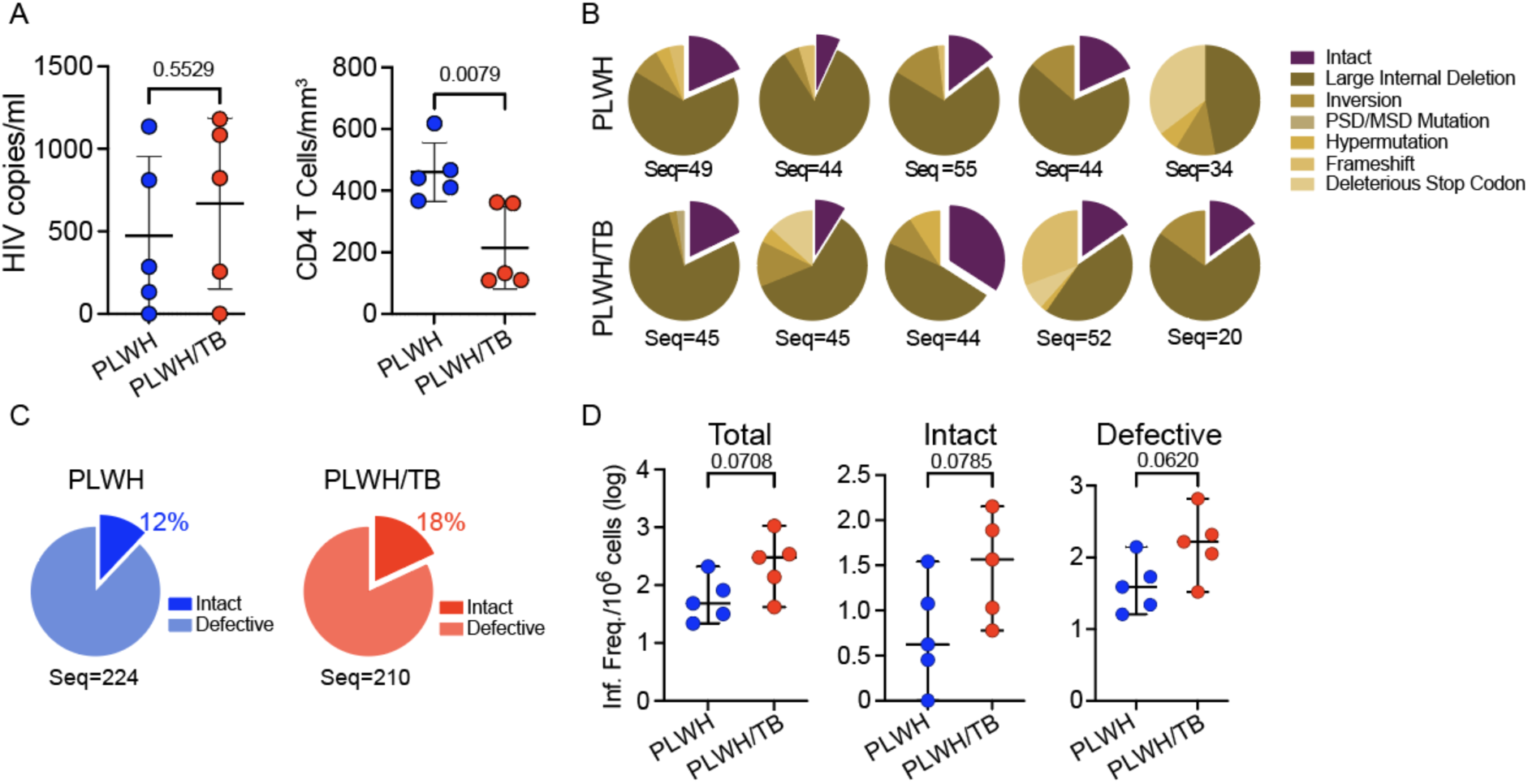
HIV-1 proviral genetic landscape in people living with HIV-1 and TB. (A) Plasma HIV-1 RNA copies per ml and CD4+ T cell counts in five PLWH/TB (red) and five PLWH (blue). (B) Pie charts showing the proportion of genetically intact and defective proviral HIV-1 sequences for each donor. (C) Averaged proportion of genetically intact and defective proviral HIV-1 sequences across groups. The total number of sequences analyzed per group is shown below each pie chart. (D) Infection frequency of total, genetically intact, and genetically defective proviruses per 10⁶ CD4+ T cells. Frequencies were calculated based on the proportion of CD4+ T cells in each sample and log-transformed. Each data point represents a single donor. Bars represent the mean ± SD. Statistical significance was determined by unpaired t-test, with p < 0.05 considered significant.

To characterize the HIV-1 proviral genetic landscape in both groups, we employed the full-length individual proviral sequencing assay (FLIPS), obtaining a total of 434 HIV-1 DNA sequences from the 10 participants. As reported before in PLWH, the majority of sequences were genetically defective (65.9%-100%), with internal deletions being the most frequent defect (47.1%-94.9%) (Figure 1B) (1). Genetically intact HIV-1 sequences were identified in nine of the ten participants (Figure 1B). The proportion of genetically intact proviruses in PLWH/TB was 18%, and 12% in PLWH (Figure 1C). When infection frequency was calculated, we observed that in PLWH/TB the frequency of infected cells per million CD4+ T cells tended to be higher for total (p=0.0708), genetically defective (p=0.0620), and genetically intact (p=0.0785) HIV-1 DNA sequences (Figure 1D). These results suggest that concurrent TB may increase the frequency of cells infected with HIV-1.

### The proportion of genetically intact HIV-1 provirus is a significantly higher at the site of the coinfection

The majority of studies on HIV-1 infection in PLWH/TB, including analysis of the HIV-1 reservoir, have been conducted using blood samples. However, we and others have shown that at the site of the coinfection the TB-associated microenvironment can influence HIV-1 pathogenesis (24-27, 29, 30). For instance, in people who developed tuberculous pleural effusion (TB-PE), the number of HIV-1 RNA copies is higher in this fluid when compared with matched plasma (Figure 2A, Supplementary Figure 1) (24-26). This result led us to investigate the impact of the TB-associated microenvironment on the HIV-1 genetic landscape at the site of the coinfection. To this end, we accessed PBMCs and pleural-effusion-derived mononuclear cells (PEMCs) samples from three PLWH/TB and performed FLIPS. One of these participants was under ART at the time of sample collection, with undetectable plasma viral load, but a relatively high level of HIV-1 RNA copies in the TB-PE (Figure 2A, Table 2). In contrast, two of these participants were untreated for HIV-1 at the time of sampling (Figure 2A, Table 2). Interestingly, the proportion of genetically intact HIV-1 DNA sequences in PEMCs was higher when compared to PBMCs across all three participants (0%-25% PBMCs; 17.8%-61.7% PEMCs) (Figure 2B). In participant 122, who was virally suppressed on ART, no intact provirus was found in the four sequences obtained from the blood. However, 17.8% of the proviral sequences were genetically intact in cells from the pleural space. Importantly, when HIV-1 infection frequency was calculated, we observed a significantly higher frequency of cells infected with genetically intact HIV-1 provirus in TB-PE when compared with peripheral blood (fold change = 41.3, p=0.0307) (Figure 2C). Our data suggest that the TB-associated microenvironment may promote the persistence of genetically intact HIV-1 at the site of the coinfection in PLWH/TB.

**Figure 2.**
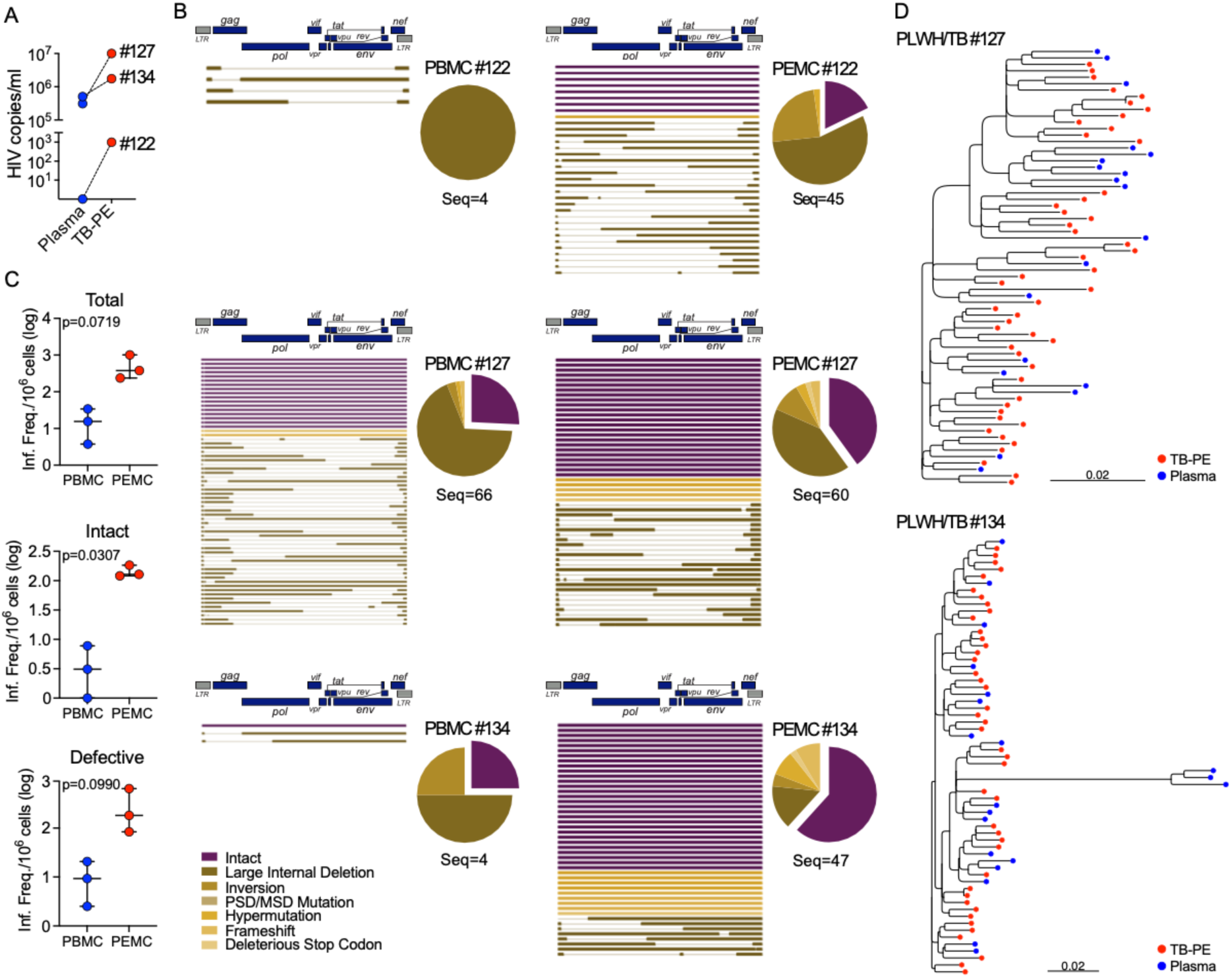
The site of HIV-1/TB coinfection is enriched in HIV-1 DNA and RNA. (A) HIV-1 RNA copies per ml in plasma (blue) and TB-PE (red) from three PLWH/TB. (B) Pie charts showing the proportion of genetically intact and defective proviral HIV-1 sequences derived from PBMCs and PEMCs for each donor. Horizontal lines adjacent to pie charts represent individual contig sequences aligned to HXB2. (C) Log-transformed infection frequency of total, genetically intact, and genetically defective proviruses per million cells. Each data point represents a single donor. Bars represent the median with 95% CI. Statistical significance was determined by paired t-test, with p < 0.05 considered significant. (D) Phylogenetic tree of HIV-1 RNA sequences derived from plasma (blue) and TB-PE (red).

### Genomic HIV-1 RNA derived from TB-PE and plasma are not genetically distinct

Given the increased frequency of genetically intact HIV-1 and elevated HIV-1 RNA levels in TB-PE compared with paired blood (Figure 2A-C and Supplementary Figure 1), we investigated whether the pleural space represents a ‘sanctuary’ site for enhanced viral replication. If this compartment was isolated from the peripheral circulation, genetic divergence between virus in the TB-PE and the plasma would be expected (26). To test this, we sequenced HIV-1 RNA derived from plasma and TB-PE samples from two untreated PLWH/TB (Donors 127 and 134). Maximum likelihood phylogenetic analysis of HIV-1 RNA sequences revealed no compartmental clustering patterns between the HIV-1 RNA derived from the pleural space and the plasma (Figure 2D). The lack of genetic divergence between viruses from both compartments indicates that the higher levels of cells infected with genetically intact HIV-1 and HIV-1 RNA copies in the pleural space cannot be explained by an isolated ‘sanctuary’ promoting local and enhanced viral replication.

### Effector memory CD4+ T cells are preferentially infected by HIV-1 in TB-PE

To determine what cell type are preferentially infected during active TB, we performed single-cell RNA sequencing (scRNAseq) on PEMCs from one PLWH/TB (donor 127) using the BD Rhapsody platform. Since the cellular composition of TB-PE in PLWH/TB has not been well defined, we compared our dataset with publicly available scRNAseq data of PEMCs from TB-PE taken from participants with *Mtb* mono-infection (Figure 3A) (33). Clustering patterns were similar across datasets, suggesting comparable PEMC phenotypes and supporting the reliability of our data (Figure 3A). Cell type identification analysis of the PEMCs from the PLWH/TB showed a predominance of CD4+ and CD8+ T cells in the large primary cluster (Figure 3B), aligning with the lymphocytic predominance of TB-PE described in previous studies (34, 35). Further characterization of the CD4+ T cell cluster identified both central and effector populations as well as several T-helper (Th) subsets including Th1, Th17, Th2, Th1/17, and T follicular helper (Tfh) cells (Figure 3C). As reported in people experiencing active TB, TB-PE-derived CD4+ T cell populations were dominated by central memory and Th1 subsets (36, 37).

**Figure 3.**
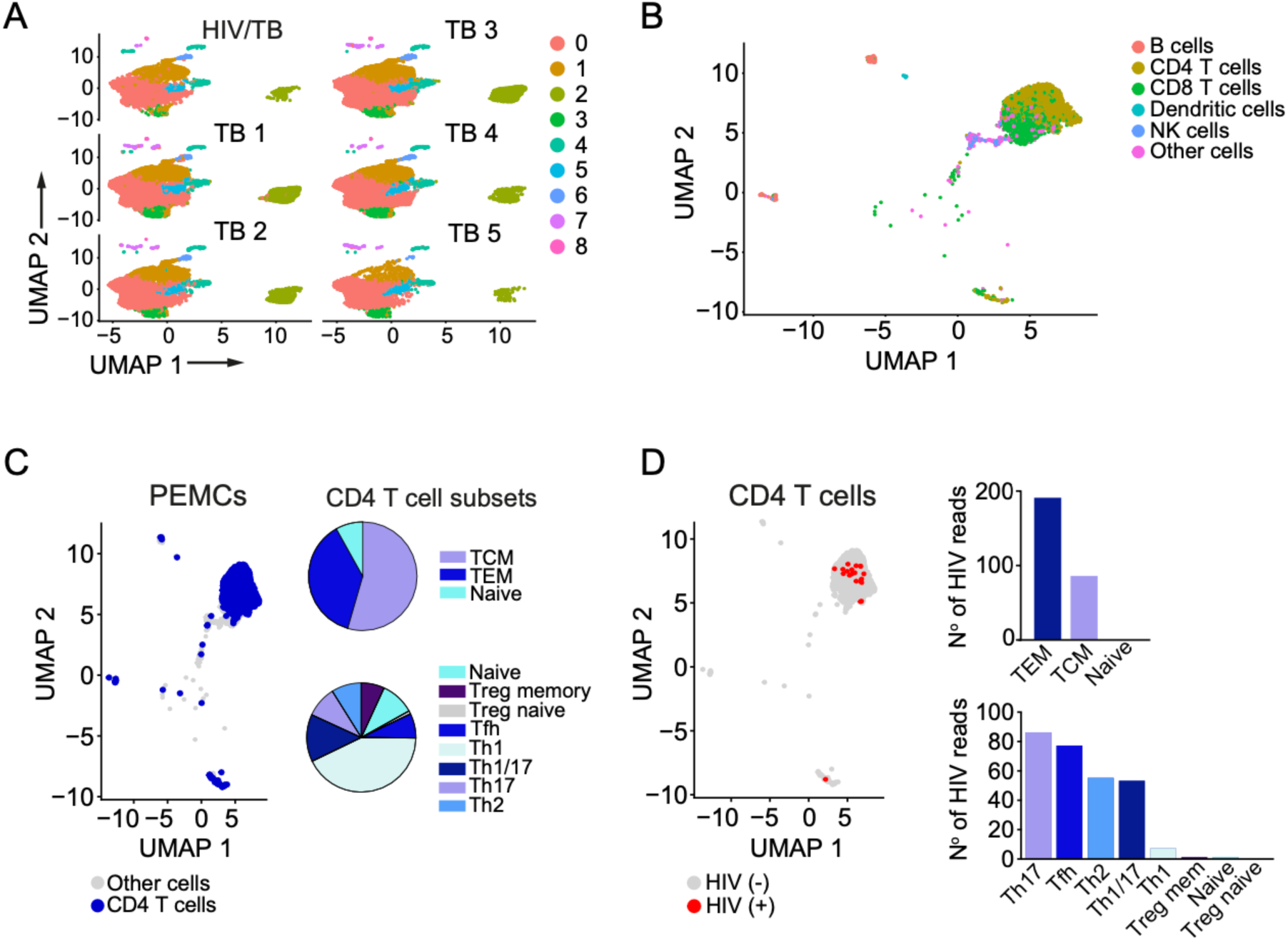
Single-cell RNA sequencing of TB-PE reveals preferential infection of effector memory CD4+ T cells. (A) UMAP of PEMCs from one PLWH/TB sample analyzed in-house compared with PEMCs from five TB patient datasets reported by Cai et al. (B) UMAP of immune cell populations from PEMCs of a PLWH/TB sample, with each cell color-coded by major immune cell type. (C) UMAP showing clusters of CD4+ T cells within PEMCs. Pie charts indicate the proportions of Tcm, Tem, and naïve CD4+ T cells (top), and Treg memory, Treg naïve, and Th subsets (bottom). (D) UMAP highlighting cells expressing HIV-1 transcripts (red). Bar graphs show the distribution of HIV-1 reads among Tem, Tcm, and naïve cells (top), and Treg memory, Treg naïve, and Th subsets (bottom). Tcm, central memory; Tem, effector memory; Treg, regulatory T cells; Tfh, follicular helper T cells; Th1, T helper 1; Th2, T helper 2; Th17, T helper 17; Th1/17; T helper 1/17.

Taking advantage of the proviral sequences previously obtained from donor 127, we generated a consensus HIV-1 sequence that was used as a reference to identify HIV-1 transcripts in our sample. HIV-1 transcripts were detected across all CD4+ T cell subsets except for the naïve Treg population (Figure 3D). In agreement with previous studies, HIV-1 transcripts were more prevalent in effector memory CD4+ T cells (Tem) when compared to central memory (Tcm) and naïve subsets (Figure 3D) (38). Interestingly, our analysis also revealed that HIV-1 transcripts were enriched within the Th17 subset, followed by Tfh, Th2, and Th1/17 cells respectively (Figure 3D). This result contrasts with previous reports showing that blood-derived Th1 CD4+ T cells are enriched for HIV-1 RNA in PLWH, but aligns with recent studies demonstrating preferential infection of Th17 CD4+ T cells in tissue (39-45). Overall, our findings suggest that the TB-associated microenvironment may promote the accumulation of preferential targets for HIV-1 infection at the site of the coinfection.

### The majority of TB-PE-derived CD8+ T cells do not express canonical markers of tissue-residency and exhibit an activated/exhausted phenotype

Our results showing an increase in the proportion of genetically intact HIV-1-infected cells in TB-PE (Figure 2) suggest reduced immune pressure at the site of coinfection. Since CD8+ T cells are critical for HIV-1 control, we investigated whether CD8+ T cells derived from TB-PE are phenotypically dysfunctional. It has been previously reported that CD8+ T cells in the airways of PLWH exhibit a tissue-resident phenotype, defined by the expression of CD103 (46), which is associated with reduced cytotoxic potential (47, 48). To determine whether TB-PE is enriched for tissue-resident CD8+ T cells, we analyzed the phenotype of PEMC-derived CD8+ T cells from two PLWH/TB (donors 127 and 134) using a 24-colour flow cytometry panel. Integration of both datasets, followed by unbiased clustering identified nine distinct clusters (Figure 4A-D). CD103 expression was restricted to clusters 4, 8 and 9, indicating that although a fraction of the TB-PE-derived CD8+ T cells are potentially tissue resident, they do not represent the majority (Figure 4C and D). CD69, another marker associated with tissue residency, was highly expressed in cluster 8 (Figure 4A and D). However, this cluster also displayed high levels of the activation markers CD38 and HLA-DR, suggesting that in this context CD69 primarily reflects activation rather than tissue residency (Figure 4D). In addition to cluster 8, CD38 and HLA-DR co-expression was also observed across clusters 6 and 7, together with the exhaustion markers PD-1 and TIGIT (Figure 4D). PD-1 and TIGIT were also detected in cluster 5. CD39, which is associated with a terminally exhausted phenotype (49), was co-expressed with CD38 and HLA-DR in cluster 7 (Figure 4D). This data suggests the presence of an expanded pool of chronically activated CD8+ T cells with an exhausted phenotype. Of note, CCR5 was widely expressed across most clusters, consistent with migration of effector cells from the blood into the inflamed pleural space (Figure 4D) (33). Clusters 1, 2, and 4 were characterized by the expression of CCR7, the costimulatory molecules CD27 and CD28, as well as the homeostatic cytokine receptor CD127, indicating a less differentiated phenotype within these subsets (Figure 4D).

**Figure 4.**
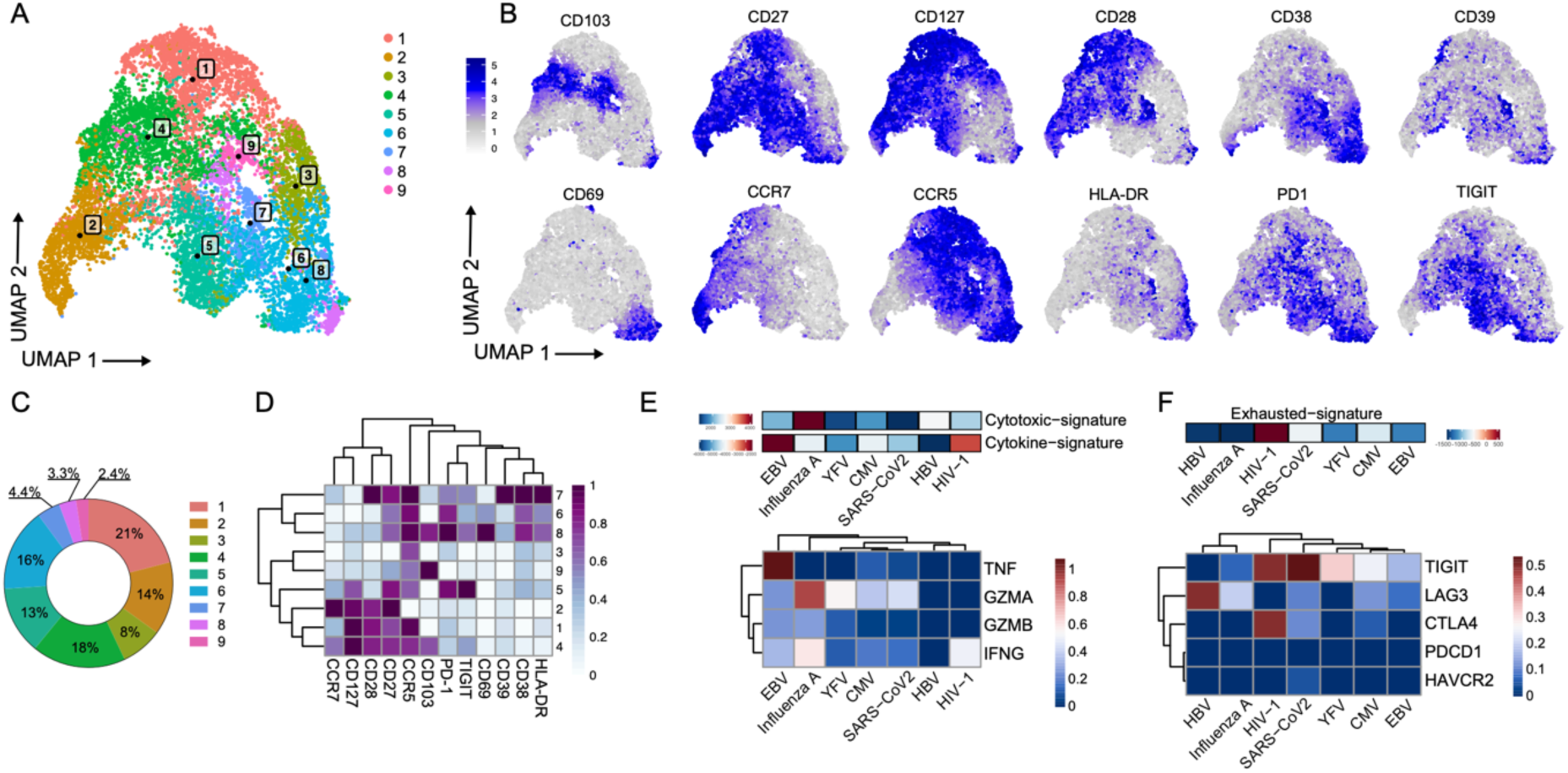
Immunophenotype of TB-PE-derived CD8+ T cells. (A) UMAP plot showing unbiased clustering of CD8+ T cells based on the immune markers CD103, CD69, CD27, CCR7, CD127, CCR5, CD28, HLA-DR, CD38, 287 PD1, CD39, TIGIT. (B) UMAP plots of expression levels of selected immune markers in TB-PE-derived CD8+ T cells. (C) Proportions of CD8+ T cell clusters identified by UMAP analysis. (D) Heatmap of average expression of 12 variably expressed markers across clusters. (E) scRNA/TCR-seq analysis of TB-PE-derived CD8+ T cells from one PLWH/TB, showing TCR specificities for viral antigens. Cytokine and cytotoxic signatures from scRNAseq data are shown (top), with a heatmap of TNF, IFNG, GZMA, and GZMB expression (bottom). (F) Exhaustion signature from scRNAseq data (top), with a heatmap showing relative expression of exhaustion-associated genes TIGIT, LAG3, CTLA4, PDCD1, and HAVCR2 (bottom).

Together, these data show that TB-PE-derived CD8+ T cells are phenotypically diverse, and that the majority do not exhibit a tissue-resident phenotype. Moreover, we identified CD8+ T cells displaying features of chronic activation and exhaustion, consistent with an inflammatory microenvironment and suggesting impaired antiviral function at the site of coinfection.

### HIV-1-specific CD8+ T cells exhibit a functionally exhausted phenotype in TB-PE

Our phenotypic analysis suggested the presence of chronically activated and exhausted CD8+ T cells at the site of the coinfection. However, this analysis did not allow us to determine whether the CD8+ T cells exhibiting this phenotype were specific for HIV-1. To address this, we have performed scRNAseq combined with single-cell T cell receptor sequencing (scTCRseq) on PEMCs from Donor 127. Based on publicly available databases (50), we identified CD8+ T cell-derived TCRs with specificity for viral antigens, including HIV-1, Epstein-Barr virus (EBV), Hepatitis B Virus (HBV), Influenza A, SARS-CoV-2, yellow fever virus (YFV), and Cytomegalovirus (CMV) (Figures 4E and F). We then analyzed the transcriptomic profile of these cells and compared the cytokine and cytotoxic signatures across the CD8+ T cells with different specificities to assess effector functionality. HIV-1-specific CD8+ T cells showed a strong cytokine signature relative to CD8+ T cells with other specificities, mainly driven by the expression of *IFNG,* indicating recent contact with the antigen (Figure 4E). However, their cytotoxic signature was considerably lower than that of other virus-specific CD8+ T cells (Figure 4E). This was further supported by reduced expression of the cytotoxic genes granzyme A (*GZMA*) and granzyme B (*GZMB*) (Figure 4E). To further understand this result, we also assessed the relative exhausted signature of HIV-1-specific CD8+ T cells. Notably, HIV-1-specific CD8+ T cells exhibited a strong exhausted signature, driven by high expression levels of *TIGIT* and *CTLA4* (Figure 4F). Collectively, these results indicate that HIV-1-specific CD8+ T cells are functionally exhausted, potentially contributing to reduced immune pressure at the site of the coinfection.

### The TB-associated microenvironment inhibits effector CD8+ T cell functionality

Exhaustion of CD8+ T cells is a well-established hallmark of HIV-1 pathogenesis (7, 8). Therefore, our finding that HIV-1-specific CD8+ T cells in TB-PE are functionally exhausted is not unique to coinfection with *Mtb*. To better understand the specific contribution of the TB-associated microenvironment to CD8+ T cell functionality, we developed an *in vitro* model using pooled acellular fractions of TB-PE from individuals with active pleural TB (Figure 5A) (30). In this model, we stimulated CD8+ T cells isolated from healthy donors with anti-CD3/CD28 antibodies to mimic antigenic stimulation. Stimulated and unstimulated CD8+ T cells were cultured in the presence or absence of TB-PE to determine whether the inflammatory TB-associated microenvironment affected the CD8+ T cell effector response (Figure 5A). We first performed bulk RNA sequencing to characterize the transcriptomic profile of cells stimulated with anti-CD3/CD28 antibodies exposed to TB-PE. Differential gene expression (DGE) analysis identified 228 upregulated genes and 237 downregulated genes between controls and cells exposed to TB-PE (supplementary Figure 2, Figure 5B and C). Gene ontology pathway analysis revealed that cell pathways and genes associated with T cell effector functions were significantly downmodulated by TB-PE (Figures 5D-F). Moreover, cell pathways related to chemotaxis were upregulated, reflective of the inflammatory nature of the TB-associated microenvironment (Figure 5E). We further validated these results by measuring the cell-surface protein expression of the activation markers CD69, CD25, and HLA-DR. Consistent with the transcriptomic profile shaped by TB-PE, the presence of TB-PE impaired the upregulation of these activation markers upon T cell stimulation (Figure 5G). To test whether the TB-PE-mediated inhibition of T cell activation is a common feature of pleural TB, we tested additional pooled and individual TB-PE samples and pleural effusion from individuals with heart failure (HF-PE). Unlike HF-PE, both the additional pool and the individual TB-PE samples reproduced the inhibitory effect (Figure 5H). These findings suggest that the impairment of CD8+ T cell activation is a consistent feature of TB-PE.

**Figure 5.**
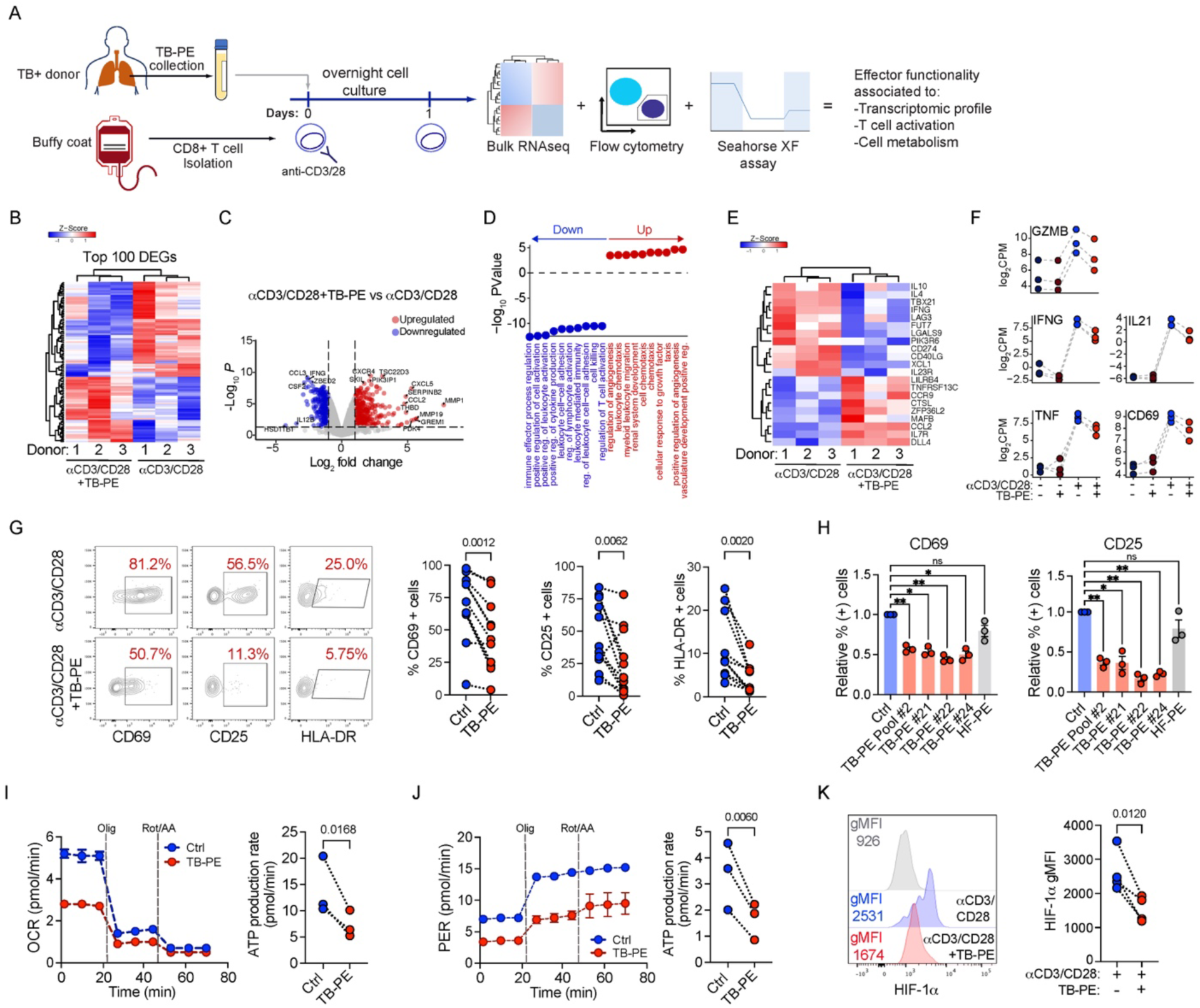
TB-PE treatment downmodulates T cell effector functionality. (A) Schematic of the experimental model used to study the effect of TB-PE on CD8+ T cell functionality. (B) Heatmap showing hierarchical clustering of the top 100 variable genes between TB-PE–treated and control CD8+ T cells stimulated with anti-CD3/CD28 antibodies. (C) Volcano plot of differentially expressed genes (DEGs). (D) Gene set enrichment analysis (GSEA) of DEGs (p < 0.05 and absolute logFC > 1) comparing TB-PE–treated and control activated CD8+ T cells. Dot plots illustrate the top 10 upregulated and downmodulated enriched GO biological process terms. (E) Heatmap of normalized expression of selected genes involved in T cell activation. (F) Gene expression levels of GZMB, IFNG, TNF, IL21, and CD69 across conditions. (G) Primary CD8+ T cells were stimulated with anti-CD3/CD28 antibodies in the presence of TB-PE; cells activated in the absence of PE served as controls. Expression of activation markers CD69, CD25, and HLA-DR was measured by flow cytometry after 24h (CD69, CD25) or 48h (HLA-DR). Representative flow plots (left) and quantification of activated cells (right) are shown. (H) Proportions of CD69 (left) or CD25 (right) positive cells stimulated in the presence of an additional TB-PE pool, three independent TB-PE samples (red), or HF-PE (grey). Values are relative to the non-PE control condition (blue). Data are shown as mean ± SD. Each dot represents an independent donor. Statistical significance was determined by paired two-tailed t-test or one-way ANOVA followed by Tukey’s HSD post-test. p ≤ 0.05, p ≤ 0.01. (I) Oxidative phosphorylation measured as oxygen consumption rate (OCR, left) and mitochondrial-dependent ATP production (right). (J) Glycolysis measured as proton efflux rate (PER, left) and glycolysis-dependent ATP production (right). (K) HIF-1α expression in activated CD8+ T cells determined by flow cytometry. Each dot represents an independent donor. Statistical significance was determined by paired two-tailed t-test or one-way ANOVA followed by Tukey’s HSD post-test. p ≤ 0.05, p ≤ 0.01.

### TB-PE treatment downmodulates oxidative phosphorylation and glycolysis

Efficient T cell responses require increased glycolysis and oxidative phosphorylation (OXPHOS)(51). We therefore assessed the metabolic profile of CD8+ T cells exposed to TB-PE using the Seahorse cell flux analyzer. Oxygen consumption rate (OCR) and the proton efflux rate (PER) were measured in CD8+ T cells as indicators of OXPHOS and glycolysis, respectively (Figures 5I and J). The intracellular rate of ATP production derived from glycolysis (glycoATP) or OXPHOS (mitoATP) was also quantified (Figures 5I and J). Consistent with the inhibition of T cell activation, both OXPHOS and glycolysis levels were significantly lower in TB-PE-treated cells when compared to controls (Figures 5I and J).

To further characterize the impact of the TB-associated microenvironment on T cell metabolism and effector functions, we also measured the expression of hypoxic inducible factor 1 alpha (HIF-1α). This transcription factor is a master regulator of glucose cell metabolism and necessary for an efficient T cell effector response (52). Cells exposed to TB-PE exhibited reduced expression of HIF-1α (Figure 5K). Taken together, these data indicate that the TB-associated microenvironment impairs CD8+ T cell activation and effector functionality upon antigenic stimulation

### HIV-1-specific CD8+ T cell responses are inhibited by TB-PE treatment

Our results demonstrate that the TB-associated microenvironment impairs the effector functionality of CD8+ T cells after antigen stimulation. We wondered whether TB-PE treatment can also inhibit the antiviral function and HIV-1-specific effector responses mediated by CD8+ T cells derived from PLWH. To address this question, we stimulated PLWH-derived PBMCs with a pool of HIV-1 peptides to expand HIV-1-specific CD8+ T cells for 14 days. After expansion, cells were restimulated with the HIV-1 peptides in the presence or absence of TB-PE, to induce the expression of the degranulation markers CD107a/b and the effector cytokines IFN-γ and TNF-α (Figure 6A). Importantly, cells treated with TB-PE exhibited significantly lower expression of CD107a/b, IFN-γ, and TNF-α when compared to cells expanded without TB-PE (Figures 6B and C). Of note, we observed no significant differences between stimulated cells and cells stimulated in the presence of HF-PE (Figure 6B and C). These results were further confirmed when cells from healthy donors were stimulated with a pool of peptides for common viruses (CMV, EBV and influenza; CEF peptide pool), in the presence or absence of TB-PE (Supplementary Figure 3). Overall, these data demonstrate that the TB-associated microenvironment inhibits the ability of CD8+ T cells to mount an efficient antiviral effector response.

**Figure 6.**
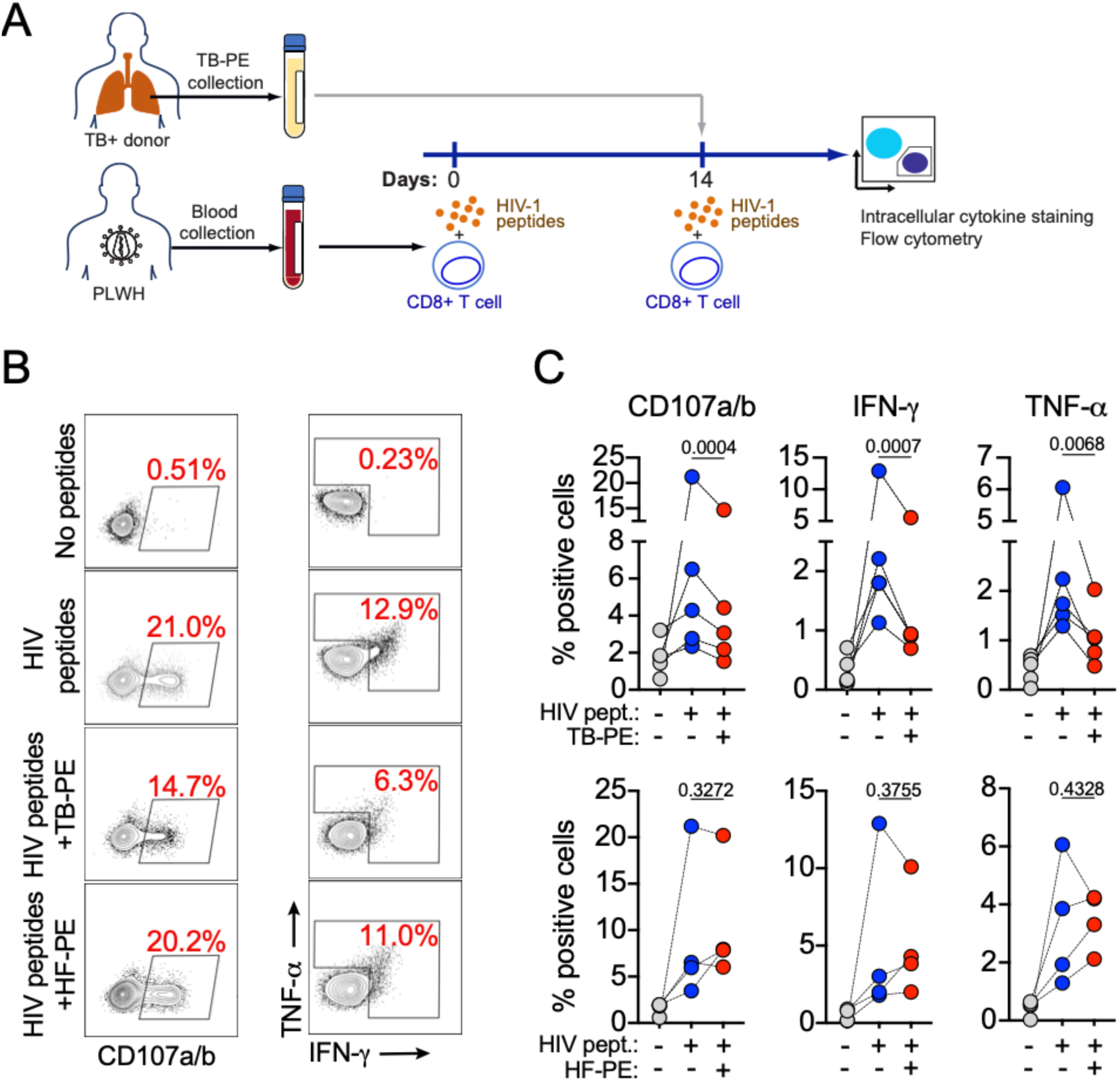
The effector anti-HIV-1 responses of CD8+ T cells are inhibited by TB-PE. (A) Schematic of the experimental model used to study the effect of TB-PE on anti-HIV-1 CD8+ T cell effector responses. (B) Representative flow cytometry of PLWH-derived CD8+ T cells expressing CD107a/b, TNF-α and IFN-γ as markers for effector response upon restimulation with HIV-1 peptides in the presence or absence of TB-PE. Cells exposed to no peptides were used as an experimental control. Cells exposed to HF-PE were used as a control to determine the specific effect of TB-PE. (C) Quantification of expression of CD107a/b, TNF-α and IFN-γ in PLWH-derived CD8+ T cells exposed to TB-PE (n=5; upper) or HF-PE (n=4; bottom). Each data point represents a single donor. Statistical significance was determined by paired t-test, with p < 0.05 considered significant.

### TB-PE prevents HIV-1-specific CD8+ T cells from acquiring cytotoxic function

Considering that TB-PE inhibits efficient TCR-dependent T cell activation, we hypothesized that the TB-associated microenvironment could also impair the differentiation of CD8+ T cells into a cytotoxic phenotype upon antigen encounter, leading to inefficient killing of HIV-1-infected CD4+ T cells. To assess this, PBMCs from PLWH were expanded with HIV-1 peptides in the presence or absence of TB-PE for 14 days. After expansion, CD8+ T cells were cocultured with autologous CD4+ T cells superinfected with HIV-1 *in vitro* and, in the absence of TB-PE (Figure 7A). Of note, at the time of coculture, the number of CD8+ T cells was normalized based on the expression of CD107a/b, ensuring that equal numbers of cytotoxic CD8+ T cells were included across conditions (Figure 7A). As expected, HIV-1-specific CD8+ T cells expanded with peptides alone significatively reduced the number of HIV-1-infected CD4+ T cells when compared to the control culture of CD4+ T cells (Figure 7B). In contrast, HIV-1-specific CD8+ T cells expanded in the presence of TB-PE were unable to eliminate HIV-1-infected CD4+ T cells when compared to CD8+ T cells expanded with peptides alone (Figure 7B). This finding was further confirmed when cells from healthy donors were expanded with the CEF peptide pool, in the presence or absence of TB-PE and cocultured with target CD4+ T cells pulsed with CEF peptides (Supplementary Figure 4). These results indicate that the TB-associated microenvironment hampers the differentiation of HIV-1-specific CD8+ T cells into fully cytotoxic effectors upon antigen exposure.

**Figure 7.**
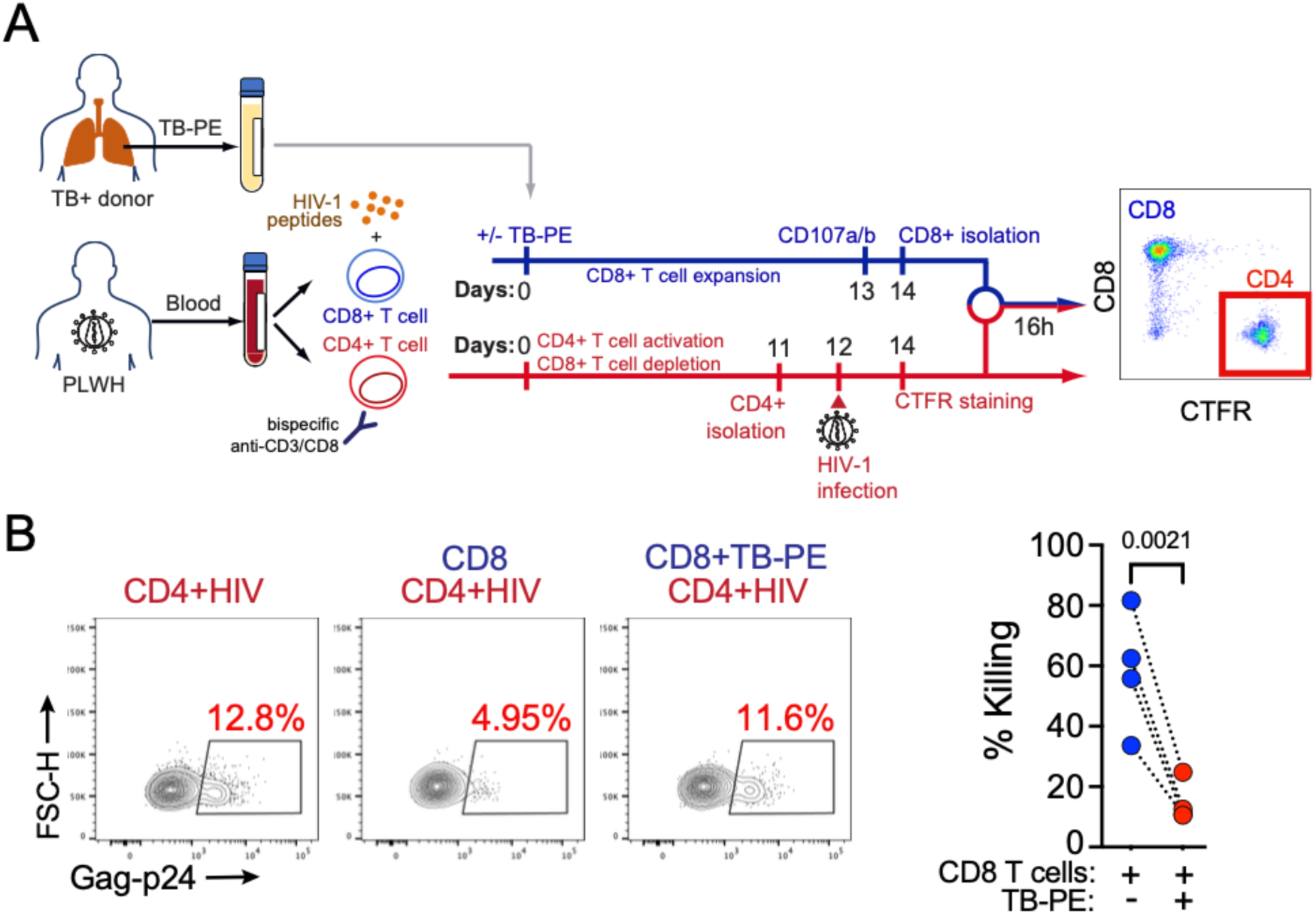
TB-PE inhibits the cytotoxic function of HIV-1-specific CD8+ T cells. (A) Schematic of the experimental model used to study the effect of TB-PE on HIV-1-specific CD8+ T cell cytotoxicity. (B) Representative flow cytometry plots showing p24 levels in HIV-1-infected CD4+ T cells cocultured with CD8+ T cells expanded in the presence or absence of TB-PE. As a control, CD4+ T cells were cultured without CD8+ T cells. Quantification of HIV-1-infected cell killing was performed using samples from four PLWH. Each data point represents a single donor. Statistical significance was determined by paired t-test, with p < 0.05 considered significant.

Taken together, our results suggest that the higher proportion of genetically intact HIV-1-infected cells observed at the site of coinfection may result from impaired CD8+ T cell-mediated control of HIV-1, driven by reduced effector differentiation and function.

## Discussion

While extensive clinical evidence highlights the synergistic interaction between HIV-1 and *Mtb*, leading to worsened outcomes for PLWH/TB, most research has focused on how HIV-1 infection increases the risk and severity of TB. By contrast, little is known about how TB disease can promote HIV-1 persistence. In this study, we addressed this gap by characterizing both the HIV-1 genetic landscape and CD8+ T cell–mediated antiviral immunity at the site of coinfection in PLWH/TB. Our findings provide two key insights: (i) the site of HIV-1/*Mtb* coinfection, represented by TB-PE, is enriched in genetically intact and replication-competent HIV-1; and (ii) the local microenvironment suppresses effective HIV-1-specific CD8+ T cell responses. To our knowledge, this is the first study characterizing the HIV-1 proviral landscape using near–full-length HIV-1 sequencing while simultaneously assessing the impact of the TB-associated microenvironment on CD8+ T cell-mediated control of HIV-1 in PLWH/TB. Taken together, these results position the TB-associated microenvironment as a modulator of antiviral responses, leading to reduced immune pressure on the concurrent HIV-1 infection and creating a permissive niche for the persistence of genetically intact, replication-competent virus.

Although previous studies have reported higher levels of HIV-1 DNA in blood-derived cells from PLWH/TB, the proviral genetic landscape in this population remains poorly characterized. A recent study showed that circulating CD4+ T cells from PLWH with a history of active pulmonary TB harbored higher levels of intact provirus compared to those without a history of active TB (17). In line with these findings, we observed in our cohort that both defective and intact HIV-1 infection frequencies tended to be higher in PLWH/TB than in PLWH without TB. Although these differences did not reach statistical significance, likely due to the small sample size, they are consistent with prior reports showing elevated levels of HIV-1 DNA in PLWH/TB (17, 19). Importantly, these observations align independently of cohort characteristics, suggesting that regardless of treatment history for HIV-1 or TB, coinfection promotes the persistence of HIV-1. This underscores the need for a deeper understanding of the mechanisms driving this virological outcome, with the goal of improving strategies to prevent and treat HIV-1 infection in PLWH/TB.

Despite increasing research on HIV-1/*Mtb* coinfection, characterizing the anatomic sites of coinfection remains challenging. Animal models, particularly non-human primates (NHPs), have provided important insights into pathogen dynamics within the lung and granuloma (53-55). However, these studies mainly assess the impact of HIV-1 infection on TB outcomes and do not fully recapitulate the pathology observed in humans. Thus, models using human samples are essential to better understand coinfection in PLWH/TB. In this study, we used TB-PE as a model of the site of coinfection to characterize the inflammatory environment shaped by *Mtb*. Consistent with previous reports, we observed higher HIV-1 RNA levels in pleural fluid compared with plasma (24, 26). Whether this increase reflects an enrichment of cells carrying genetically intact and replication-competent virus at the site of the coinfection has not been investigated. Using near–full-length sequencing assays developed in our laboratory, we provide, to our knowledge, the first characterization of HIV-1 proviruses in the pleural compartment, revealing an enrichment of genetically intact sequences within the proviral pool and a higher frequency of cells harboring intact proviruses in TB-PE. This enrichment of genetically intact HIV-1 at the coinfection site could be explained by two potential mechanisms: (i) enhanced local viral replication promoted by the inflammatory TB microenvironment and/or (ii) reduced antiviral immune pressure within this environment. Given the high mutation rate of HIV-1, an isolated compartment would be expected to display genetic divergence between viruses. However, our phylogenetic analysis of paired plasma and TB-PE samples did not reveal genetic divergence, suggesting that the elevated levels of HIV-1-infected cells at the coinfection site in these individuals cannot be attributed to an isolated “sanctuary” supporting independent viral replication. Previous reports sequencing the HIV-1 envelope region did identify genetic differences between PE and plasma but also demonstrated viral trafficking between the two compartments (26).

Although we previously showed that the TB-associated microenvironment can promote HIV-1 spread in macrophages, we also demonstrated that TB-PE does not increase HIV-1 replication in CD4+ T cells (27, 29, 30). Of note, T-cell lymphocytes are the dominant cell type in TB-PE (21, 23, 56). These findings suggest that the mechanisms by which the TB-associated inflammatory microenvironment promotes HIV-1 persistence in CD4+ T cells within the TB-PE are likely driven by reduced immune pressure rather than enhanced viral replication.

Substantial efforts in the HIV-1 field have aimed to identify CD4+ T-cell subsets preferentially infected by HIV-1. A growing body of evidence has highlighted Tem CD4+ T cells as a key target, being more susceptible to infection and enriched in genetically intact proviruses (38, 57). Within the spectrum of CD4+ T cell subsets, preferential HIV-1 infection has also been reported in T helper subsets such as Th1, Th2, Th17, Th1/17, and Tfh (42-44, 58). More recently, single-cell technologies have enabled a deeper characterization of HIV-1 target cells in both blood and tissue (39, 40, 59, 60). Th1 CD4+ T cells with cytotoxic features have been identified as a persistent source of HIV-1 in blood, whereas Th17 and Th1/17 subsets appear to be preferentially infected in tissues (40, 59, 60). Despite these advances, it remains unclear which cells are preferentially infected at the site of HIV-1/*Mtb* coinfection. In the current study, using scRNA-seq in TB-PE, we mapped HIV-1 transcripts across pleural CD4+ T cell subsets. Consistent with previous studies in PLWH, we found that Tem CD4+ T cells were enriched in HIV-1 transcripts (39, 40, 59). Interestingly, infected cells were primarily concentrated within the Th17, Tfh, Th2, and Th1/17 subsets. Notably, although Th1 CD4+ T cells represented the dominant subset in TB-PE, this population contained relatively few HIV-1 transcripts, which contrasts with the high-level of HIV-1 transcripts produced by Th1 CD4+ T cells in the blood (40). Finally, unlike prior reports, we did not detect evidence of productive infection in Treg (61), which may reflect differences in the methods used to phenotype TB-PE-derived cells.

As mentioned above, the enrichment of genetically intact HIV-1 at the site of the coinfection could result from either enhanced local viral replication or reduced antiviral immune pressure. Since our results suggest that the pleural compartment is not a privileged site of viral replication in CD4+ T cells, we next assessed whether CD8+ T cell-mediated control of HIV-1 is compromised at this site. Recent evidence indicates that tissue-resident CD8+ T cells display reduced cytotoxic capacity, contributing to HIV-1 persistence in tissues such as lymph nodes (9, 10). In contrast to immunological studies of bronchoalveolar lavage-derived CD8+ T cells from PLWH, we observed that only a fraction of the CD8+ T cell population in TB-PE exhibited a tissue-resident phenotype (47, 62). These differences may reflect the distinct cellular composition of the lung parenchyma versus the exudative nature of pleural TB. Notably, a subset of TB-PE-derived CD8+ T cells displayed markers of chronic activation and exhaustion. Importantly, HIV-1-specific CD8+ T cells in TB-PE exhibited an exhausted phenotype with diminished cytotoxic potential, which likely contributes to inefficient HIV-1 control at this site. Because CD8+ T cell exhaustion is a well-known feature of HIV-1 infection, independent of *Mtb* coinfection (63), we sought to define the specific impact of the TB-associated microenvironment. Using our *in vitro* model, we demonstrated that the inflammatory pleural environment impairs CD8+ T cell effector functionality, leading to suboptimal antiviral responses. Consistent with this, CD8+ T cells exposed to TB-PE were unable to eliminate HIV-1-infected cells, supporting the idea that the higher proportion of genetically intact HIV-1 in PLWH/TB may arise from reduced immune pressure. Although pleural TB is a transient condition, the establishment of a pool of cells enriched in genetically intact proviruses may have long-term implications for HIV-1 reservoir persistence. Indeed, one participant in our study had undetectable plasma viremia but displayed relatively high HIV-1 RNA levels and intact proviruses within TB-PE, likely reflecting both limited drug penetration into this compartment and impaired CD8+ T cell-mediated control. These findings underscore the importance of understanding immune functionality at the site of HIV-1/*Mtb* coinfection.

Our work provides new insights into the immune mechanisms that may contribute to worsened virological outcomes in PLWH/TB. It underscores the need for further exploration of therapeutic interventions aimed at minimizing the impact of active TB on the HIV-1 reservoir. These findings also have important implications for the HIV-1 cure field as we provide evidence that, at the site of coinfection, impaired HIV-1-specific CD8+ T cell responses may contribute to the accumulation of genetically intact, replication-competent proviruses in PLWH/TB, representing an additional challenge for cure-focused interventions in this population.

## Limitations

In this study, we provide a comprehensive characterization of the virological and immunological features of the HIV-1/*Mtb* coinfection site, modeled by TB-PE. However, several limitations should be acknowledged. First, access to these rare samples was limited, and our genetic comparison between proviruses derived from blood and TB-PE included only three PLWH/TB. Despite the small sample size, our findings were consistent, showing a high proportion of genetically intact HIV-1 in this compartment, including in one participant on effective ART. Second, our single-cell transcriptomic analysis was performed in only one participant, and matched blood samples were not available. Nevertheless, comparison with publicly available datasets revealed similar clustering patterns, supporting the validity of our observations. Finally, to determine the specificity of CD8+ T cell-derived TCRs, we relied on public databases. While these repositories are inherently limited to reported TCR sequences and thus may not capture the full spectrum of antigen-specific TCRs present in our sample, this limitation is comparable to studies using tetramers, where analyses are restricted to predefined specificities.

## Methods

### Ethical Statement

This research was carried out in accordance with the Declaration of Helsinki (64). This study was approved by the institutional review boards at the Hospital F. J Muñiz (NIN-2601-19) and the Academia Nacional de Medicina de Buenos Aires (12554/17/X); Facultad de Medicina, Universidad de Buenos Aires (Buenos Aires, Argentina); and the Western Sydney Local Health District which includes the Westmead Institute for Medical Research (2022/PID00100 - 2022/ETH00092). Buffy coats from healthy donors were obtained from the Australian Red Cross Blood Service, Sydney, Australia. Written informed consent was obtained from all participants in this study.

### Participant cohort and clinical samples

To genetically characterise HIV-1 proviral sequences, PBMCs isolated by Ficoll density gradient centrifugation were collected from eight PLWH/TB and five PLWH and cryopreserved at -80°C. The samples were collected at the Hospital F.J. Muniz, Buenos Aires, Argentina. The diagnosis of active TB was bacteriologically confirmed by either positive acid-fast bacilli smear, Ziehl-Nielsen smear, or *Mtb* culture. Relevant clinical data of each participant are described in Tables 1 and 2. Three of the PLWH/TB had been diagnosed with pleural TB disease requiring therapeutic thoracentesis at the Hospital F. J. Muniz in Buenos Aires, Argentina. Therapeutically aspirated tuberculous pleural effusions were collected as described below and stored at -80°C until use. To investigate the effect of TB-PE on HIV-1-specific CD8+ T cell responses, PBMCs from five PLWH on ART recruited through the Phaedra/Core01 AIEDRP protocol (Australia) were collected. Relevant clinical data of each participant are described in Supplementary Table 2.

### Pleural effusion samples

Pleural effusions (PE) were obtained by therapeutic thoracentesis at the Hospital F.J Muniz (Buenos Aires, Argentina) from three PLWH/TB (Table 2) and 13 HIV-1-negative people with PE (Supplementary Table 1). The diagnosis of TB-PE was based on a positive Ziehl-Nielsen stain or Lowestein-Jensen culture from pleural effusion and/or histopathology of pleural biopsy. The diagnosis was confirmed by an *Mtb*-induced IFN-γ response and a positive adenosine-deaminase test. Exclusion criteria included the presence of concurrent communicable or non-communicable (cancer, diabetes, or steroid therapy) conditions, and multi-drug resistant *Mtb*. For the HIV-1-negative donors, exclusion criteria also included a positive HIV-1 test. PE were collected in heparin tubes and centrifuged at 300g for 10 minutes at room temperature without brake. Cells from PLWH/TB were cryopreserved and stored at -80°C. The cell-free supernatant was transferred into new plastic tubes and further centrifuged at 12,000g for 10 minutes at room temperature and aliquots were stored at -80°C.

### Pleural effusion pools

PE from recruited HIV-1-negative participants were divided into two pools depending on aetiology. The first pool was comprised of samples from nine participants diagnosed with tuberculous pleural effusion (TB-PE) and the second comprised of four patients with heart-failure-induced transudates (HF-PE). Pools were prepared by combining equal amounts of PE derived from each donor. The prepared pools were decomplemented at 56°C for 30 minutes and filtered through a 0.22μm mesh to remove residual bacteria or debris.

For PE-treated experimental conditions, 20% v/v PE was added to the cell culture. This concentration of PE was previously optimised by our group (29, 65-67).

### Primary CD8+ T cell isolation from healthy donors

PBMCs were obtained by Ficoll-Hypaque (Sigma Aldrich) density gradient centrifugation from buffy coats of healthy donors. CD8+ primary T cells were isolated and purified from PBMCs by negative selection using the Human CD8+ T cell isolation kit (Miltenyi Biotec).

### Full-Length Individual Proviral Sequencing (FLIPS)

Near-full-length HIV-1 proviral sequences were obtained using the FLIPS assay as previously described (68, 69).

### Identification of genetically intact and defective HIV-1 proviruses

Proviruses were considered intact if they passed the criteria of the Proviral Sequence Annotation and Intactness test (https://psd.cancer.gov/tools/tool_index.php) and met the following criteria: (1) all open reading frames (ORFs) had an intact start codon, (2) regulatory/accessory protein ORFs were intact (no insertions or deletions>5% of the ORF length, stop codons, or frameshift mutations), (3) possessed a cryptic major splice donor (MSD) 4 nucleotides downstream of MSD 1 in the event MSD 1 was mutated.

Defective HIV-1 proviruses were categorised as full-length (>8800bp), deleted (<8800bp), or genomically rearranged (containing an inversion or repeated region). After categorising deleted and genomically rearranged proviruses, full-length proviruses were sequentially characterised as hypermutant using the Los Alamos National Laboratory Hypermutation tool (https://www.HIV-1.lanl.gov/content/sequence/HYPERMUT/hypermutv3.html), containing an MSD or packaging signal deletion (HXB2 position 695-810), containing a frameshift mutation in any ORF, or containing a stop codon in any ORF.

ORF intactness was assessed on the presence of an intact start codon, an absence of insertions/deletions >5% of the length of the ORF, premature stop codons, and frameshift mutations. Splice donor and acceptor sites were not assessed in this study due to the genetic diversity of splice sites between participants, previous evidence showing that HIV-1 protein expression can still occur in proviruses with defective splice sites, and evidence that HIV-1 can utilise cryptic splice sites when traditional splice sites are defective (70-72).

Sequences were classified as identical if they were 100% identical according to the Los Alamos National Laboratory ElimDupes tool (https://www.hiv.lanl.gov/content/sequence/elimdupesv2/elimdupes.html).

### Plasma-derived RNA using long-range sequencing assay (PRLS)

Genomic HIV-1 RNA derived from the plasma and PE were sequenced using the plasma-derived RNA using long-range sequencing assay (PRLS) as previously described (73). This assay enables near–full-length sequencing of individual plasma/TB-PE-derived HIV-1 RNA genomes spanning *gag* to the 3′ LTR (*gag*-3′; ∼8.3 kb). Due to reduced efficiency resulting from incomplete reverse transcription in samples with low viral loads, a modified version of the PRLS assay targeting the HIV-1 *integrase* gene to the 3′ LTR (*int*-3′; ∼5 kb) was used when indicated (73). For the plasma sample of donor 127, we applied the *gag-3’* assay. For all other samples the *int-3’* assay was used.

### Phylogenetic Analysis

Genomic HIV-1 RNA sequences were trimmed to the *integrase*-3’ LTR region for phylogenetic analysis between sequences derived from the plasma and the PE. Maximum likelihood trees were constructed for matched plasma and PE-derived sequences. Sequences with an internal deletion (>100bp) and sequences identified as hypermutant by the Los Alamos National Laboratory Hypermut tool (https://www.hiv.lanl.gov/content/sequence/HYPERMUT/hypermutv3.html) were excluded from this analysis. Sequences were visualised using maximum likelihood phylogenetic trees constructed using FastTree version 2.2.0 (74) using the generalised time-reversible model. Branch support values were calculated using 1000 bootstrap replicates and resulting phylogenetic trees were visualised and annotated using ggTree (75) to identify clustering patterns between plasma and PE-derived sequences.

### High-parameter flow cytometry

To characterize the cellular composition of TB-PE the following antibodies were included in a high-parameter flow cytometry panel: CD28-BUV395 (CD28.8, BD Biosciences), CD8-BUV496 (SK1, BD Biosciences), HLADQ-BUV563 (TU169, BD Biosciences), CD161-BUV615 (HP-3G10, BD Biosciences), CD154-BUV737 (TRAP1, BD Biosciences), CD4-BUV805 (OKT4, BD Biosciences), CCR7-BV421 (2-LI-A, BD Biosciences), CD39-BV480 (TU66, BD Biosciences), HLADR-BV605 (G46-6, BD Biosciences), PD1-BV711 (EH12.1, BD Biosciences), CD27-BV786 (LI27, BD Biosciences), CCR5-VioB515 (REA245, Miltenyi Biotec), CD3-NovaFluorB610 (UCHT1, Thermo Fisher Scientific), CD69-BB700 (FN50, BD Biosciences), TIGIT-PE (A15153G, BioLegend), CD103-PEVio615 (REA803, Miltenyi Biotec), CD127-PECY7 (R34.34, Thermo Fisher Scientific), and CD38-APCFire810 (HIT2, BioLegend).

PEMCs from two donors (2.5 x 10^6^ cells) were stained with BD Horizon™ Fixable Viability Stain 440UV (1:2000, BD Biosciences) in the dark for 30 minutes at 4°C followed by a wash with FACS buffer (1% foetal calf serum [v/v], 2 mM ethylenediaminetetraacetic acid [EDTA], 0.1% sodium azide [w/v] in PBS). Cells were then incubated in the dark at room temperature for 10 minutes with 3.2µL Oligoblock (Sigma-Aldrich) and 5µL human AB serum to block non-specific binding. Cells were then stained with a master mix of antibodies for surface chemokine receptors containing 10µL brilliant stain buffer (BD Biosciences) for 15 minutes in the dark at room temperature. The remaining surface antibodies were prepared as a master mix containing 10 µL brilliant stain buffer (BD Biosciences) and incubated with cells in the dark for 30 minutes at room temperature. Cells were then washed twice with FACS buffer and fixed in Cytofix buffer (BD Biosciences) for 20 minutes at 4°C. Cells were then washed with FACS buffer and acquired on the BD FACSymphony. Data was recorded using BD FACSDiva.

Data were processed using dimensionality reduction (Uniform Manifold Approximation and Projection; UMAP) and clustering using FlowSOM (76).

### Single-cell RNA and T-cell receptor sequencing library

Single-cell suspensions of PEMCs were prepared for the BD Rhapsody single cell sequencing pipeline according to manufacturer’s instructions. 12,000 PEMCs were processed using the BD Rhapsody system and RNA and TCR libraries were generated per cell using the TCR/BCR Full Length, mRNA Whole Transcriptome Analysis (WTA), and Sample Tag kit (BD Biosciences). Libraries were sequences at 20,000 reads per cell for WTA and TCR libraries using the Novaseq platform (Illumina).

### Single-cell RNA and T-cell receptor sequencing analysis

Initial data analysis utilised the BD Seven Bridges desktop app and standard pipelines. Further analysis was performed using the R packages Seurat (77) and scRepertoire (78) for differential gene expression analysis, TCR repertoire identification, and data visualisation. Pathway analysis was conducted with the R package Escape (79). Cell type identification was performed using the SingleR (80) and CellDex (80) packages, with the associated reference libraries including the Monaco (81) reference for CD8+ T cell subset identification, the Novershtern (82) reference to identify central and effector memory cell subsets, and the Database of Immune Cell Expression (DICE) (83, 84) reference to identify CD4+ T cell subsets. HIV-1-infected cells were identified by using the HIV-1 consensus sequences previously obtained through the FLIPS assay for this participant as a reference to identify HIV-1 transcripts via the Seven Bridges pipeline. TCR sequences for cells specific for HIV-1, cytomegalovirus, Epstein-Barr virus, Influenza, and SARS-Cov2 were obtained from the VDJdb database (50). Single-cell RNAseq data from PEMCs from participants with pleural TB alone were obtained from publicly available datasets (accession numbers HRA000910 and HRA00036) and used for comparative analysis.

### Bulk RNA sequencing library

Primary CD8+ T cells were obtained from healthy donors as described above. A subset of these cells were activated using anti-CD3/CD28 antibodies (Miltenyi Biotec) in the presence or absence of TB-PE. Separately, we cultured un-activated cells separated into TB-PE treated and untreated conditions. After 20h of stimulation, the cells were collected to perform Bulk RNAseq. RNA from CD8+ T cells (2x10^6^ cells per condition) was extracted using the RNeasy Plus Mini kit (QIAGEN). Total extracted RNA was quantified using the Qubit broad range RNA assay, and RNA quality was assessed using the Agilent 4200 Tapestation system. Library preparation was performed using 400ng of total RNA as input and the Illumina stranded mRNA Prep kit. Short read sequencing was performed on the Novaseq 6000 platform where a minimum of 20 million paired-end 50bp reads per sample were generated.

### Bulk RNA sequencing analysis

Library sequencing quality was determined using FastQC (Babraham Bioinformatics: https://www.bioinformatics.babraham.ac.uk). Illumina adapter sequence and low quality read trimming (read pair was removed if it was <20 base pairs) was performed using Trim Galore (Babraham Bioinformatics). STAR (85) was used to align reads to human genome hg38 using ENSEMBL gene annotations as a guide. Read counts data corresponding to ENSEMBL gene annotations were generated and the STAR flag – quantMode GeneCounts. Multiqc (86) was used to verify quality metrics. All analyses were performed in the R Statistical Environment (87) with the Tidyverse package (88). EdgeR was used to perform background correction and to normalise count data by library size (89). Differentially expressed gene analysis was determined using the QLFtest (90)(Benjamini-Hochberg Multiple Testing Correction p<0.05). Gene lists were functionally annotated using the Gene Ontology Biological Process (GO:BP) pathways (adjusted p value <0.05) using the clusterProfiler package (91).

### Glycolysis and oxidative phosphorylation rate measurement

CD8+ T cells were purified by magnetic isolation and stimulated with anti-CD3/CD28 antibodies (Miltenyi Biotec) in the presence or absence of TB-PE. After 24h of stimulation, cells were cultured in RPMI-based Seahorse medium (Agilent) supplemented with 10mM D-glucose, 1mM pyruvate, and 2mM glutamine and plated in a Seahorse 24-well plate at a concentration of 7x10^5^ cells/well. Proton efflux rate (PER) and oxygen consumption rate (OCR) were measured using a Seahorse XFe24 analyser. ATP production rates were measured using the Seahorse XF Real Time ATP Rate Assay kit. Oligomycin, rotenone, and antimycin A were added to quantify mitochondrial or glycolysis derived ATP production rates. PER and OCR values were normalised by calculating the cellular area per condition.

### Preparation of Viral Stocks

HIV-1^NL4-3^ stocks were produced by transfection (1 μg/well) in HEK 293T cells (3 × 10^5^ cells per well in 6-well plates) using X-treme gene transfection reagent (Sigma Aldrich). Pseudotyping was achieved by co-transfecting pHEF-VSVg (400 ng/well). Supernatant was harvested at 48h and 72 h post-transfection, cleared by centrifugation at 1,500g for 10 min, and frozen at −80°C. Viral stocks were titrated by infecting primary CD4+ T cells and flow cytometry analysis of p24 expression 72h post infection.

### HIV-1 infection

Isolated CD4+T cells (obtained as described below) were cultured at a concentration of 10^6^ cells/ml in cRF10 in a 96 well U-bottom plate and incubated overnight with the corresponding HIV-1 stock at 37°C. Then, cells were washed two times with cRF10, and fresh cRF10 was added. When indicated, the proportion of HIV-1-infected cells was determined by intracellular staining of the viral protein Gag-p24 with a Mouse anti-p24-FITC antibody (clone KC57, Beckman Coulter).

### CD8+ T cell-mediated cytotoxicity assay

*Expansion of CD8+ T Cells (Effector cells).* The expansion of virus-specific CD8⁺ T cells was performed as previously described (92). Cryopreserved PBMCs from PLWH or healthy donors were thawed and rested overnight in cRF10. The following day, cells were cultured in AIM-V medium (Thermo Fisher Scientific) in the presence or absence of TB-PE for 30 minutes. After pre-incubation, PBMCs from PLWH were stimulated with 4 µg/ml Gag, Pol, and Nef peptide pools (HIV-1 peptide pool; NIH AIDS Reagent Program), while cells from healthy donors were treated with 4 µg/ml of a peptide pool derived from cytomegalovirus, Epstein-Barr virus, and influenza (CEF peptide pool; NIH AIDS Reagent Program) for 2 h at 37 °C.

Cells were washed with RF10 and cultured in cRF10 supplemented with 100 U/ml recombinant human IL-2 (BioLegend) and 10% v/v TB-PE for 12 days. Culture medium was replaced every 48-72 h. Expanded CD8+ T cells were isolated by negative selection using the Human CD8+ T cell isolation kit (Miltenyi Biotec).

The frequency of antigen-specific and cytotoxic CD8+ T cells was determined by CD107a/b staining following peptide re-stimulation, as previously described (92). The proportion of CD107a/b⁺ cytotoxic CD8+ T cells was used to normalize effector cell numbers in subsequent co-culture assays, as detailed below.

*Expansion of CD4+ T Cells (Target cells).* In parallel, CD4+ T cells were expanded from cryopreserved PBMCs from PLWH or healthy donors. Cells were cultured in cRF10 medium supplemented with 100 U/ml IL-2 and 0.5 µg/ml CD3/CD8 bispecific antibody (ARP-12277; obtained from Johnson Wong and Galit Alter through the NIH AIDS Reagent Program).

PBMCs from PLWH were cultured in the presence of antiretrovirals (100 nM efavirenz, 100 ng/ml enfuvirtide, and 30 µM raltegravir; Millipore Sigma) to prevent viral spread and reinfection. Treatment with the CD3/CD8 bispecific antibody selectively depleted CD8+ T cells, enriching for activated CD4+ T cells. On day 11, CD4+ T cells were purified by negative selection using the Human CD4+ T Cell Isolation Kit (Miltenyi Biotec).

*Effector-Target co-culture.* On day 11, purified CD4+ T cells from PLWH were infected *in vitro* with HIV-1. After three days of infection, cells were stained with 1 µM CellTrace Far Red (CTFR; Thermo Fisher Scientific) and co-cultured with autologous CD8+ T cells at a 1:1 effector:target ratio.

For healthy donors, CD4+ T cells were pulsed with the CEF peptide pool for 1 h at 37 °C to generate antigen-specific target cells. These “target” cells were stained with 0.1 µM CTFR, whereas un-pulsed “bystander” CD4⁺ T cells were stained with 1 µM CTFR. Target, bystander, and expanded CD8+ T cells were co-cultured at a 1:1:1 ratio (target:bystander:effector).

After 24 h of co-culture, CD8+ T cell-mediated killing was quantified by measuring intracellular p24 expression in HIV-1-infected targets or by calculating the ratio of bystander to target cells in CEF-pulsed assays.

### Quantification and Statistical Analysis

Experimental data was analysed using Prism (GraphPad Software). Normality of the data was tested using the Kolmogorov-Smirnov test, or the Shapiro-Wilk test in a sample-size dependent manner. Based on the results of the relevant normality test, statistical significance was tested with either a paired or unpaired t test to compare two conditions, or a one-way ANOVA followed by Tukey’s HSD post-test or Kruskal– Wallis followed by Dunn’s post-test were used for multiple comparisons. HIV-1 infection frequency per million cells was calculated using a binomial model.

## Acknowledgments

We acknowledge with gratitude the participants who donated samples for this study. Flow cytometry and RNA-seq analysis were performed at the Westmead Scientific Platforms, which is supported by the Westmead Research Hub, the Cancer Institute New South Wales, the National Health and Medical Research Council, and the Ian Potter Foundation. We would like to thank Dr. Joey Lai, Genomics Facility manager at The Westmead Institute for Medical Research, for the RNA-seq library preparation. This work was supported by the Delaney AIDS Research Enterprise to Find a Cure (1UM1AI126611-01 and 1UM1AI164560-01), the Australian National Health and Medical Research Council (APP1149990), the Agence Nationale de Recherche sur le Sida et les hépatites virales (ANRS2018-02, ECTZ 118551/118554, ECTZ 205320/305352, ANRS ECTZ103104 and ECTZ101971), Sidaction (13457), the Argentinean National Agency of Promotion of Science and Technology (PICT 2019-2019-01044 and PICT 2020-00501), Sandra and David Ansley, the Sydney Medical School Foundation, and The University of Sydney Institute for Infectious Diseases.

## Author contributions

G.D. conceptualized and designed the study. G.D. and S.C. designed and conducted the majority of *ex vivo* and *in vitro* experiments. G.D., L.B., and S.P. provided supervision, critical input, and funding support for the project. L.B., F.Q., A.K. and G.T. provided clinical samples. J.S. and S.C. performed the transcriptomic analyses. A.P.C., K.F., and E.L. assisted with HIV-1 sequencing. Y.L. and T.O. contributed to flow cytometry data analysis. F.W.v.D. and K.B. performed high-parameter flow cytometry staining. J.M.R. assisted with *in vitro* experiments. Z.V. and C.V. performed metabolic assays. S.C. and G.D. wrote the original manuscript. L.B. and S.P. supervised and revised the final version of the manuscript.

