## Supplementary material for "The tuberculosis-associated microenvironment promotes HIV-1 persistence by impairing CD8+ T cell-mediated viral control": Table 1

**Table 1. Clinical data for participants included in this study.**

| Participant ID | Sex assigned at birth | Age | Viral Load [A] (copies/ml) | CD4+ T cell count [A] (cells/μl) | Time on ART [A] (months) | HIV therapeutic regimen [A] | TB anatomic site | TB therapeutic regimen |
|---|---|---|---|---|---|---|---|---|
| 255472 | F | 37 | <50 | 410 | 3 | Data not available | n/a | n/a |
| 290480 | F | 34 | 810 | 367 | 1 | Data not available | n/a | n/a |
| 304060 | M | 26 | 1136 | 441 | 3 | TDF/FTC/EFV | n/a | n/a |
| 259876 | F | 24 | 285 | 467 | 3 | 3TC/AZT/NVP | n/a | n/a |
| D000078C | M | 31 | 133 | 618 | 2 | TDF/FTC/EFV | n/a | n/a |
| 299182 | M | 31 | <50 | 132 | 1 | TDF/FTC/EFV | Lymph node tuberculosis | 1 month HRZE |
| 294369 | M | 43 | 1180 | 109 | 96 | FTC/TDF/RAL | Lymph node tuberculosis | 2 months HRZE |
| 296604 | M | 31 | 823 | 110 | 4 | 3TC/AZT/NVP | Pleural tuberculosis | 2 months HRZE |
| 298280 | M | 38 | 257 | 359 | 2 | TDF/FTC/EFV | Lymph node tuberculosis | 5 months HRZE |
| 302124 | F | 30 | 1084 | 362 | 1 | TDF/FTC/EFV | Pulmonary tuberculosis | 2 months HRZE |

3TC, lamivudine; ABC, abacavir; ATV, atazanazir; AZT, zidovudine; COBI, cobicistat; DRV, darunavir; ECV, entecavir; EFV, efavirenz; ETR, Etravirine; EVG, elvitegravir; FPV, fosamprenavir; FTC, emtricitabine; MVC, maraviroc; NVP, nevirapine; RPV, rilpivirine; RTG, raltegravir; RTV, ritonavir; TCV, dolutegravir; TDF, tenofovir disoproxil fumarate

[A]At time of sampling.

HRZE: Isoniazid (H), Rifampicin (R), Pyrazinamide (Z) and Ethambutol (E).
