## Supplementary material for "The tuberculosis-associated microenvironment promotes HIV-1 persistence by impairing CD8+ T cell-mediated viral control": Table 2

**Table 2. Clinial data for participants included in this study.**

| Participant ID | Sex assigned at birth | Age | HIV therapeutic regimen[A] | Sample type | Viral load [A] (copies/ml) | TB therapeutic regimen[A] |
|---|---|---|---|---|---|---|
| 122 | M | 31 | FTC/TDF/DTG | Blood | 0 | None |
| | | | | Pleural Fluid | 989 | |
| 127 | M | 55 | No ART | Blood | 309909 | None |
| | | | | Pleural Fluid | >10000000 | |
| 134 | F | 47 | No ART | Blood | 507259 | None |
| | | | | Pleural Fluid | 1754399 | |

FTC, emtricitabine; TDF, tenofovir disoproxil fumarate; DTG, dolutegravir.
[A]At time of sampling.
