## Supplemental Material for "The tuberculosis-associated microenvironment promotes HIV-1 persistence by impairing CD8+ T cell-mediated viral control"

**Supplementary tables**

**Table S1. Participant Characteristics for HIV-negative donors with Pleural Effusion.**

| | **TB-PE** | **HF-PE** |
|---|---|---|
| **Age (years, range)** | 35 (19-52) | 50 (35-66) |
| **Gender, male %** | M, 78% | M, 80% |
| **Nationality, %** | Argentina, 56% Bolivia, 33% Peru, 11% | Argentina, 100% |
| **TB disease %** | Pleural, 22% Pleural and Pulmonary, 78%* | - |
| **Etiology** | *M. tuberculosis* | ND |

*Confirmed by chest X-Ray

**Table S2. Clinical data for participants included in this study**

| Participant ID | Age (years) | Sex | Time on therapy[A] (years) | Viral load[A] (copies/ml) | Antiretroviral regimen[A] |
|---|---|---|---|---|---|
| PS02017 | 29 | Male | 3 | <50 | NVP, TRU |
| PS02018 | 46 | Male | 3 | <50 | 3TC, DDI, EFV, TRU |
| PS03022 | 20 | Male | 4 | <50 | ABC, 3TC |
| PS01008 | 29 | Male | 2 | <50 | SQV |
| PS01007 | 42 | Male | 4 | <50 | ABC, 3TC |

3TC, lamivudine; ABC, abacavir; DDI, didanosine; EFV, efavirenz; NVP, nevirapine; TRU, truvada, SQV, saquinavir.

[A]At time of sampling.

**Supplementary Figures**

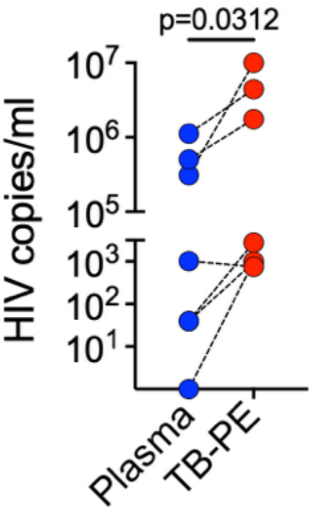


***Supplementary Figure 1. HIV viral load is higher in TB-PE.*** *HIV RNA copies per ml in plasma (blue) and tuberculous pleural effusion (red) from seven people living with HIV and pleural TB.*

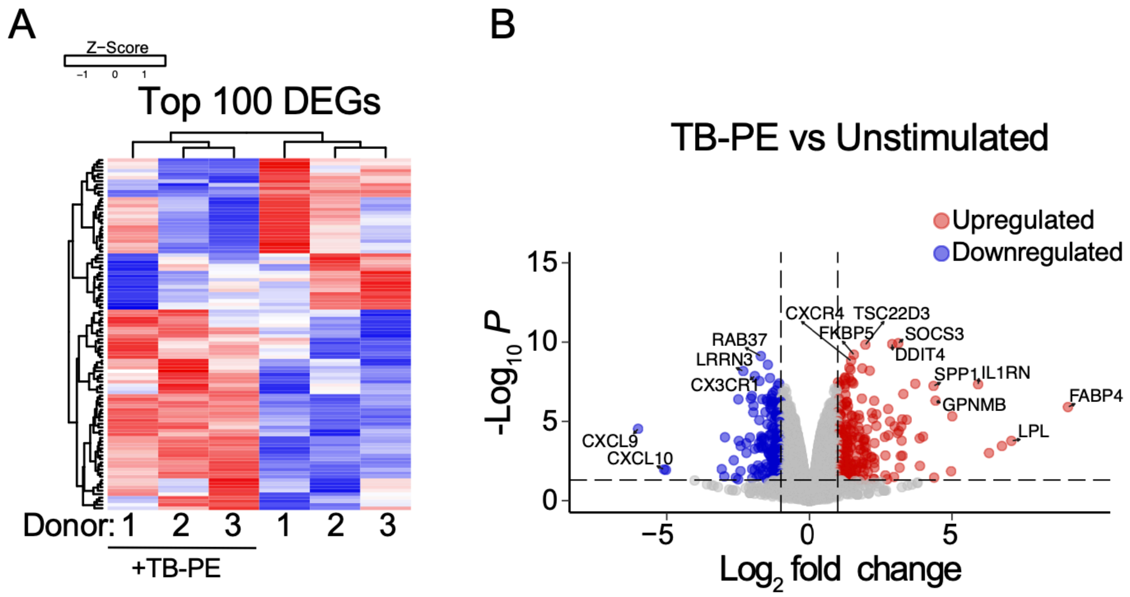


***Supplementary Figure 2. TB-PE treatment alters the transcriptome of CD8+ T cells.*** *(A) Heatmap showing hierarchical clustering of the top 100 differentially expressed genes (DEGs) between TB-PE treated and control CD8+ T cells. (B) Volcano plot of differentially expressed genes with significantly upregulated genes in red and significantly downregulated genes in blue (p<0.05).*

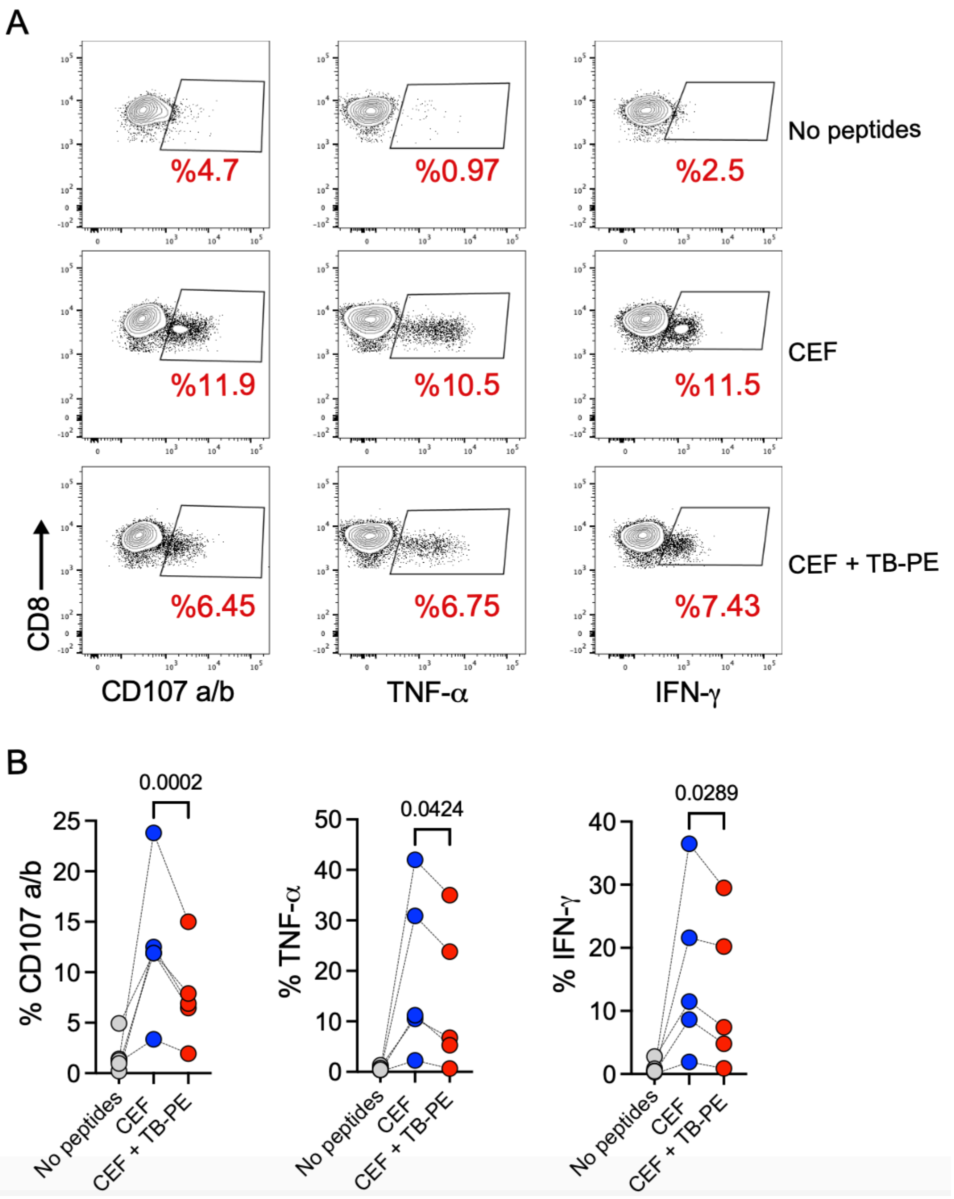


***Supplementary Figure 3. The effector antiviral responses of CD8+ T cells are inhibited by TB-PE.*** *(A) Representative flow cytometry of healthy donor-derived CD8+ T cells expressing CD107a/b, TNF-$\alpha$, and IFN-$\gamma$ as markers of effector response upon restimulation with pooled peptides from cytomegalovirus, influenza, and Epstein-Barr virus (CEF peptide pool) in the presence (red) or absence (blue) of TB-PE. Cells exposed to no peptides (grey) were used as an experimental control. (B) Quantification of the expression of CD107a/b, TNF-$\alpha$, and IFN-$\gamma$ in healthy donor-derived CD8+ T cells exposed to TB-PE (n=5). Each data point represents one donor. Statistical significance was assessed by paired t-test with $p<0.05$ considered significant.*

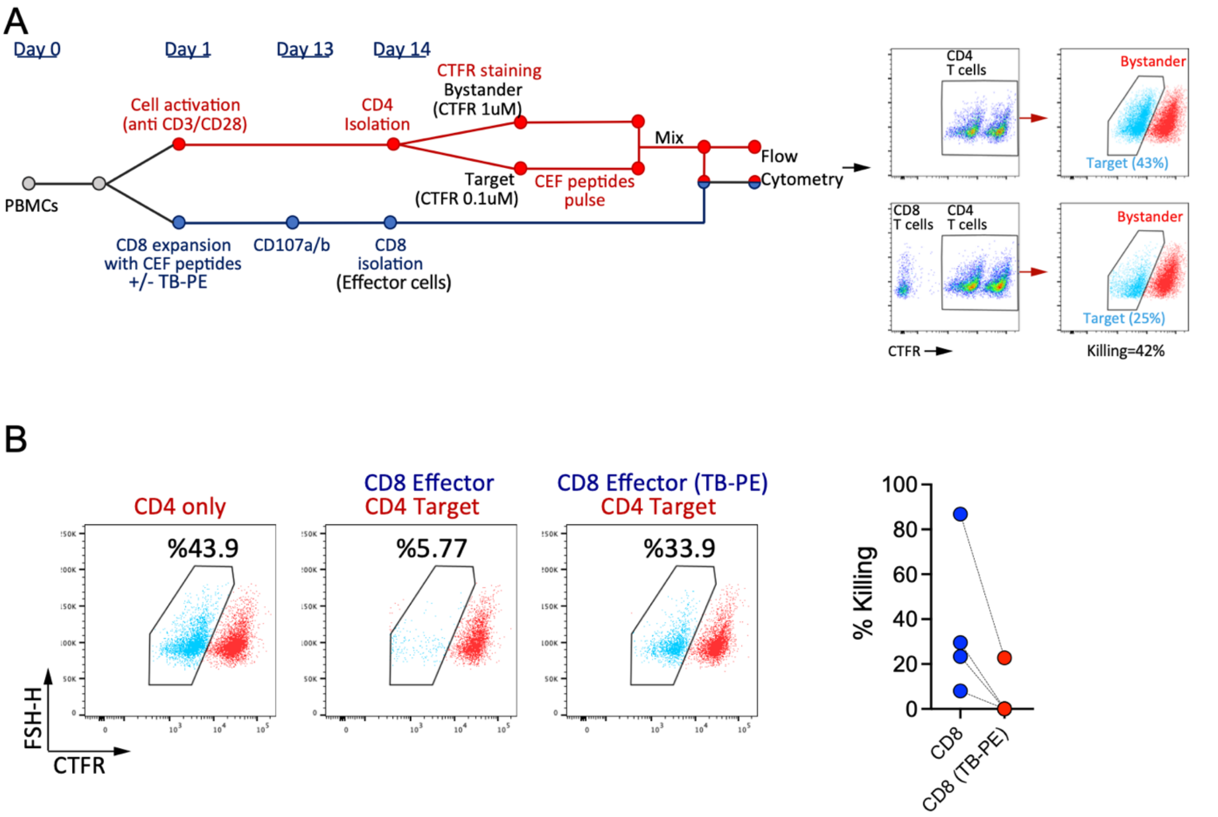


***Supplementary Figure 4. TB-PE inhibits the cytotoxic function of CD8+ T cells specific for Cytomegalovirus, Influenza, and Epstein Barr Virus.*** *(A) Schematic of the experimental model used to study the effect of TB-PE on virus-specific CD8+ T cell cytotoxicity. CD8+ T cells were expanded with a pool of peptides derived from Cytomegalovirus, Epstein Barr Virus and Influenza (CEF peptide pool) in the presence/absence of TB-PE. (B) Representative flow cytometry plots showing the proportion of target cells after coculture with CEF-specific CD8+ T cells. As a control, CD4+ T cells were cultured without CD8+ T cells (left). Quantification of CD8+ T cell-mediated killing of cells pulsed with CEF peptides (right). Each dot represents a different donor.*
